# Comparative metabolic profile of maize genotypes reveals the tolerance mechanism associated under combined stresses

**DOI:** 10.1101/2020.06.11.127928

**Authors:** Suphia Rafique

## Abstract

Drought, heat, high temperature, waterlogging, low-N stress, and salinity are the major environmental constraints that limit plant productivity. In tropical regions maize grown during the summer rainy season, and often faces irregular rains patterns, which causes drought, and waterlogging simultaneously along with low-N stress and thus affects crop growth and development. The two maize genotypes CML49 and CML100 were subjected to combination of abiotic stresses concurrently (drought and low-N / waterlogging and low-N). Metabolic profiling of leaf and roots of two genotypes was completed using GC-MS technique. The aim of study to reveal the differential response of metabolites in two maize genotypes under combination of stresses and to understand the tolerance mechanism. The results of un-targeted metabolites analysis show, the accumulated metabolites of tolerant genotype (CML49) in response to combined abiotic stresses were related to defense, antioxidants, signaling and some metabolites indirectly involved in nitrogen restoration of the maize plant. Alternatively, some metabolites of sensitive genotype (CML100) were regulated in response to defense, while other metabolites were involved in membrane disruption and also as the signaling antagonist. Therefore, the present study provides insight into the molecular mechanisms of tolerance under various stresses of maize plants, that governed on the regulation of cell wall remodeling, maintain metabolic homeostasis, defense against the pathogen, proper signaling, and restore growth under stress conditions.

## 1. Introduction

Crop faces various stresses in their life span including abiotic and biotic. Several abiotic stresses that affect crop growth, and development are drought, waterlogging, low nutrient availability, salinity, high temperature and heat. The predicted changes in the global climate has accentuate in the frequency and intensity of abiotic factors, most of the abiotic stresses occur in combination, for instant, in tropical area’s terminal drought and waterlogging occurs simultaneously during summer-rainy season. Although on many occasions the stresses are for shorter duration and relatively mild in nature, but have a significant impact on plant growth and development. Therefore, co-occurrence of abiotic stresses are more destructive to crops than their separate occurrence at different crop growth stages **(Mittler 2006; Prasad et al 2011)**. Besides, the composite responses of plants due to various concurrent stresses rely on signals, these varied and contrast signals may interact with each other to enhanced or obstruct one another **(Vile et al. 2012; Suzuki et al. 2014)**. Maize is the most important cereal, grown at the wide geographical ranges of latitude and longitude. In South Asia, particularly in tropical and subtropical environment maize is largely grown during summer-rainy season, in marginal areas which often faces extreme water availability such as waterlogging or water scarcity in a form of terminal drought. Correspondingly, co-occurrence of both the stresses often headed to the depletion in nitrogen content, thus causing low-N stress in the soil. However, studies on combined stresses by **Rizhsky et al (2004)** revealed that the response of plant to combined stresses is unique and cannot be extrapolated from the response of plant to each individual stress. Though, plants belonging to the same genus showed different molecular response in combined stresses **(Aprile et al 2013)**. Therefore, the nature of the interaction is an important aspect to fully identify the influences of combined abiotic stresses on crop plants. Among omics approach is metabolomics that has been neglected in crop improvement programs **(Kumar et al 2017)**. It has not be fully explored due to its complexity. Though, metabolites plays important role in plant metabolism that influences the plant biomass and architecture **(Turner et al 2016)**. Metabolomics study enable to identify wide variety of metabolites that gives the comprehensive view of the cellular process and physiological condition that display in a particular stress condition. Besides, the metabolites assign the functional gene and impact of the particular gene on the metabolic pathway, and that provides information of regulation as well interruptions of linked pathways **(Wen et al 2015)**. Further, metabolomics study enable to improve breeding resources through selecting the remarkable traits **(Zivy et al 2015)**. Therefore, the aim of metabolomics profiling is to compare the combined stress response in two maize inbred that differed in their stress adaptability and to unrevealed the complex molecular mechanism of tolerance at metabolomics levels in different stress conditions applied simultaneously.

## 2. Materials and Methods

### Plant Material and Growth conditions

In the present work two maize inbred (CML49 & CML100) were selected that showed different adaptability to various combine stress treatments on this basis we have categorized them as ‘tolerant and sensitive.’ The selected two inbred (CML49 and CML100) plants with 60 pots each were grown in natural conditions in greenhouse up to 30 days. 30 DAS they were subjected to various stress treatment, the plants were grown in low nitrogen, only 25% of the recommended dose of urea was applied to the treated pots. First 30 pots of each inbred were exposed to a drought stress for 10 days, while 30 control pots were supplied with full nutrient and water. After re-watering for two days normally, the same plants were exposed to waterlogging stress for up to 7 days, to maintain the water level 2-3cm above the soil surface of the pots plants were watered day and night. The roots samples from stressed and control replicates plants of each inbred (CML49 & CML100) were kept on the last day of combined stresses for extracting metabolites. Samples were quickly frozen in liquid nitrogen after removing from the plant and then kept at -80° C for further analysis.

### Sample Preparations and Metabolites extraction

The leaves and roots samples from three independent biological replicates each from control and treated maize plants (CML49 & CML100) was used for metabolite profiling. Harvested plant samples immediately frozen in liquid Nitrogen in a plastic bag (resistant to liquid N_2_). Frozen samples were stored at -80°C. The leaves and roots were homogenized separately with liquid Nitrogen in a mortar and pestle into fine powder. One gram powder of each sample was extracted three times with 5 ml of methanol at room temperature and sonicated for 15 min and centrifuged. The extract was concentrated to 1 ml by Speed Vac. The supernatant filter through Whatman filter paper (No. 4), extracts collected and stored at 4 °C for further use. The prepared extracts of both leaves and roots of the control and treated plants were subjected to gas chromatography–mass spectroscopy (GC–MS) analysis. Samples were analyzed using GC-MS for the analysis of untargeted metabolites. For derivatization, methoxylamine in pyridine (10µl) and *N*-methyl-*N*-trimethylsilyltrifuoroacetamide MSTFA (90µl) were added.

### Gas Chromatography and Mass spectrometry

The GC-MS analysis was performed with a GC-MS QP-2010 ULTRA equipped with an auto sampler (AOC-20i + s) from Shimadzu (Japan), using Equity-5 column, 30.0 m × 0.25 um × 0.25 mm for separation, and helium was used as a carrier gas at a constant flow rate of 1.0 mL/min. The 1μL sample was injected using 1/100 split-mode injections at a temperature of 260°C. The oven temperature program was initially set at 100°C and held for 2 min. The temperature was gradually increased to 250°C at a rate of 10°C/min, and 300°C at 15°C/min. Ions were generated by a 70ev electron beam at an ionization current of 2.0mA. Total ion chromatogram spectra were recorded in the mass range of 40–900 m/z at the rate of 2.5 spectra s^-1^. For metabolite identification and annotation, peaks were matched against customized reference spectrum databases including the National Institute of Standards and Technology NIST and the Wiley Registry, internal libraries and further confirmed with ChEBI (http://www.ebi.ac.uk/chebi/init.do) and ChemSpider (http://www.chemspider.com/). Data obtained was then uploaded to the web-based tool Metabo Analyst for high throughput analysis.

## 3. Results

### Comparison of metabolite levels of two inbred in combined abiotic stress condition

Metabolomics profiling using GC-MS was conducted of tolerant (CML49) and sensitive (CML100) genotypes under various combined stresses. Total 132 compounds were identified in two genotypes that includes, fatty acids, (long, medium and short chains), fatty acid derivatives, fatty aldehydes, fatty alcohols, carboxylic acids, terpenoids, glucosinolates, heterocyclic compounds, halohydrogens, alkaloids, phenolic acids, steroids and vitamins. Univariate analysis of two genotypes (CML49 and CML100) was performed by fold change analysis, t-test and volcano plots. Fold change showed the comparison between the control group metabolites with that of treated group metabolites and important features selected by t-test with threshold (p<0.05) level, the results was plotted in log 2 scale. Thus, same fold change have the same distance from zero base line. The tolerant and (CML49) and sensitive (CML100) genotypes were subjected to drought, waterlogging and low-N stresses. Tolerant genotypes contains saturated fatty acids, unsaturated fatty acids, fatty alcohols, various terpenoids, carboxylic acids, methyl esters, halocarbons, heterocyclic compounds, Strikingly, in both genotypes an overlap of Furan’s, Tiglic acid and some antioxidant compounds Tocopherol acetate, squalene and Isophytol was observed in leaves and roots. Sensitive genotype leaves shows higher accumulation of saturated and unsaturated fatty acids mostly, fatty alcohols, fatty anhydride and higher alkanes level was presented in greater amount **(Table 1)**.

**Table 1.**
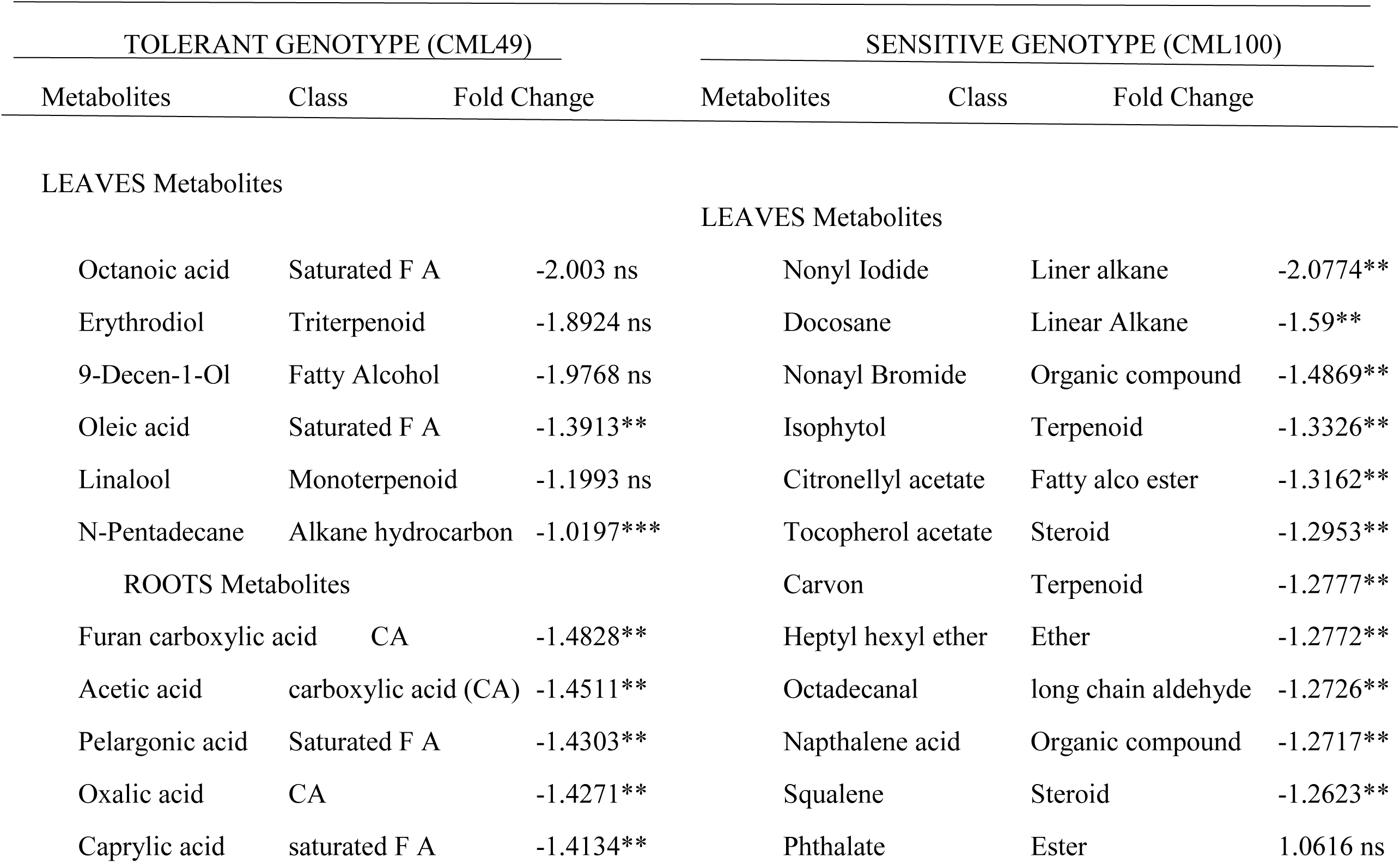
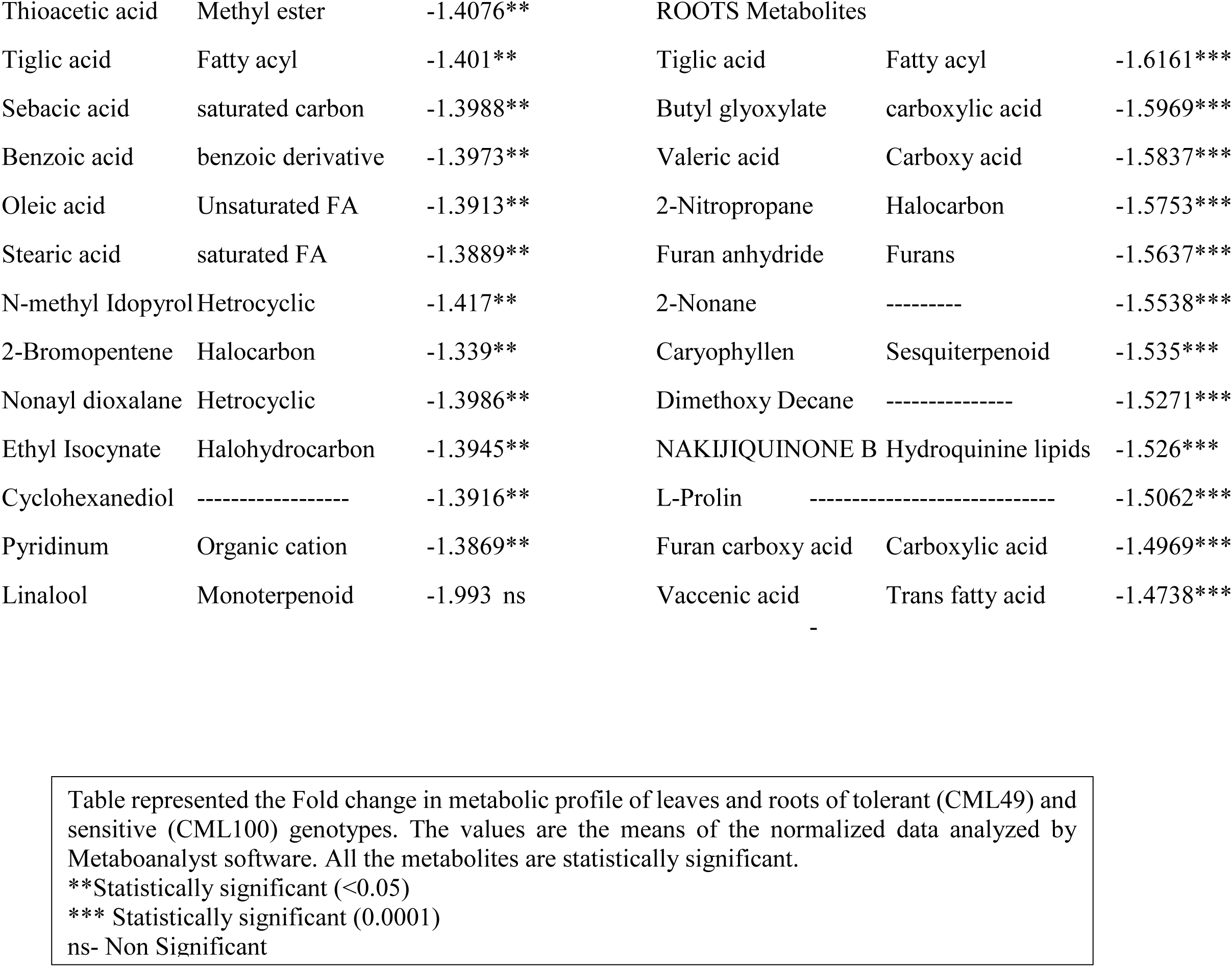
Metabolic profile of leaf and roots of tolerant and sensitive maize plants after exposure to combined abiotic stresses.

The metabolomics profile of two different genotypes suggested that the response of maize plants to various stresses applied simultaneously was shared as well as also distinctive. However, tolerant genotype roots metabolite profile display large variance in metabolites accumulation, (mostly are halohydrocarbons, heterocyclic compounds, organic cations and saturated and unsaturated fatty acids, carboxylic acids, esters) compared to roots of sensitive genotype which have less diversity in metabolite accumulation. To reduce the multivariate data complexity, to highlight the similarities and difference patterns between samples, PCA was performed for metabolites concentration data of control and treated leaves and roots of tolerant and in sensitive plants. Also, to verify the difference between the metabolic profiles of the control and treated tissues statistically and to identify the main metabolites responsible for the differences. A score scatter plots of tolerant leaf showed the good separation of metabolites between treated and control plants. In PC1 the variance is greater (70.1%) and PC2 showed variance of (19.3%). Similar trend of metabolites separation was observed of sensitive genotype leaves. The treated plants exhibited higher variances in PC1 (91.2%) and control plants variance is less PC2 (5.2%) **Fig 1a, b**.

**Fig 1.**
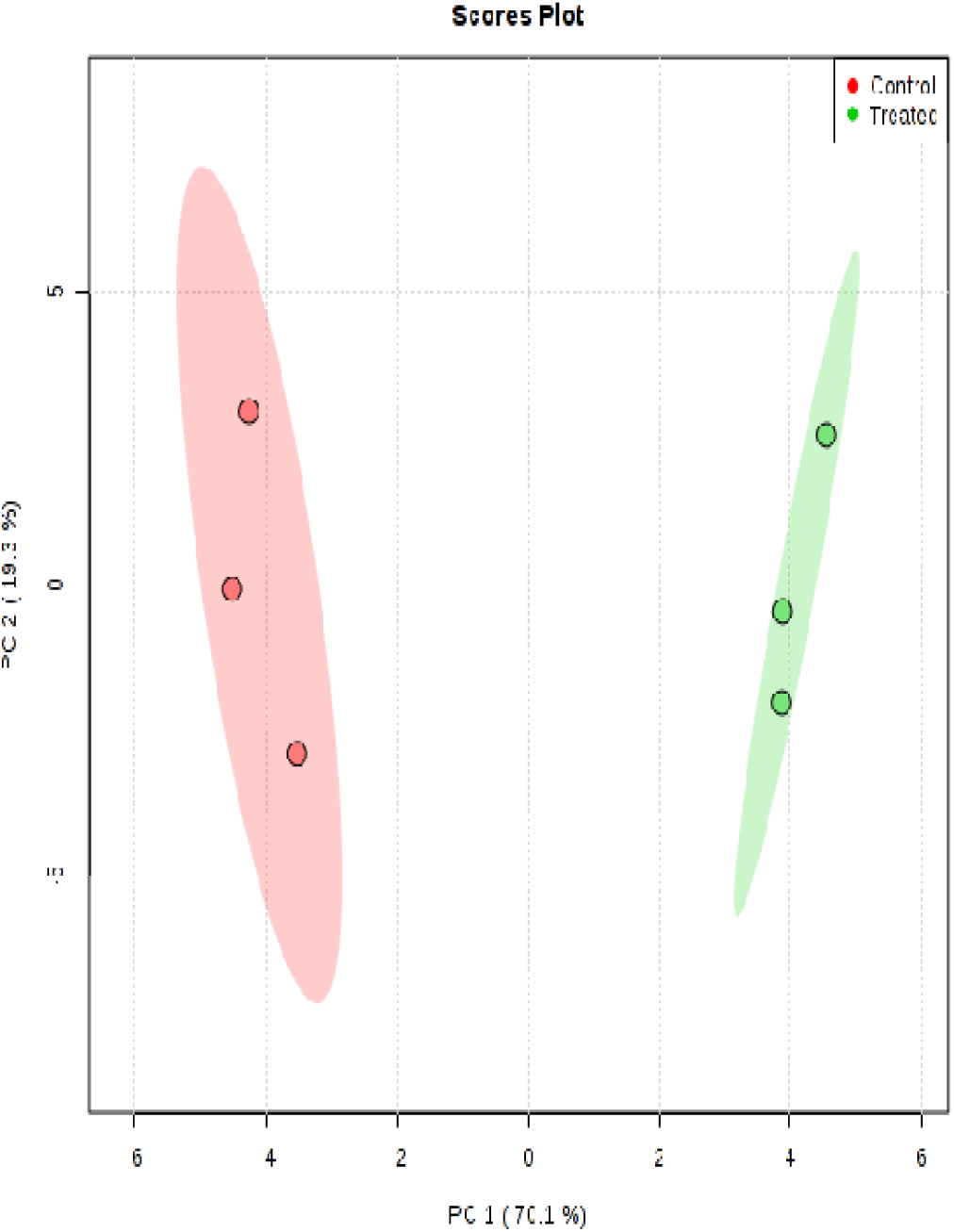
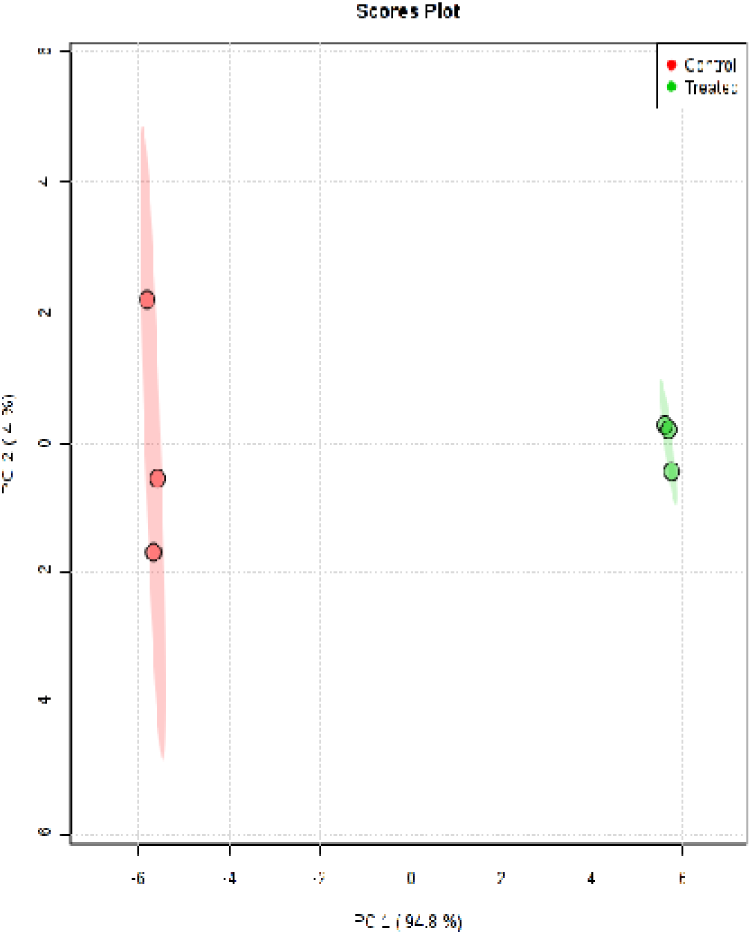
**a.** Score plot of tolerant leaf shows higher variations in metabolites of PC1 than PC2 **b.** Score plot of Sensitive leaf showing variations in metabolites of control and treated leaf. PC1 have higher variation than PC2.

However PCA of roots the trend was opposite for both genotypes, the score plot for PC1 (94.8%) and PC2 (4%) demonstrate clear separation between control (root metabolites) and treated (root metabolites) it may be due to the presence of many saturated and unsaturated fatty acids and halo-hydrocarbon in the samples. Correspondingly, sensitive (CML100) genotype PC1 (94%) and PC2 (5.3%) **Fig 2a, b**, control root metabolites were well separated and treated roots metabolites patterns shows less variance in separation because presence of similar metabolites like, 2-Furan carboxylic acid and Furan anhydride.

**Fig 2.**
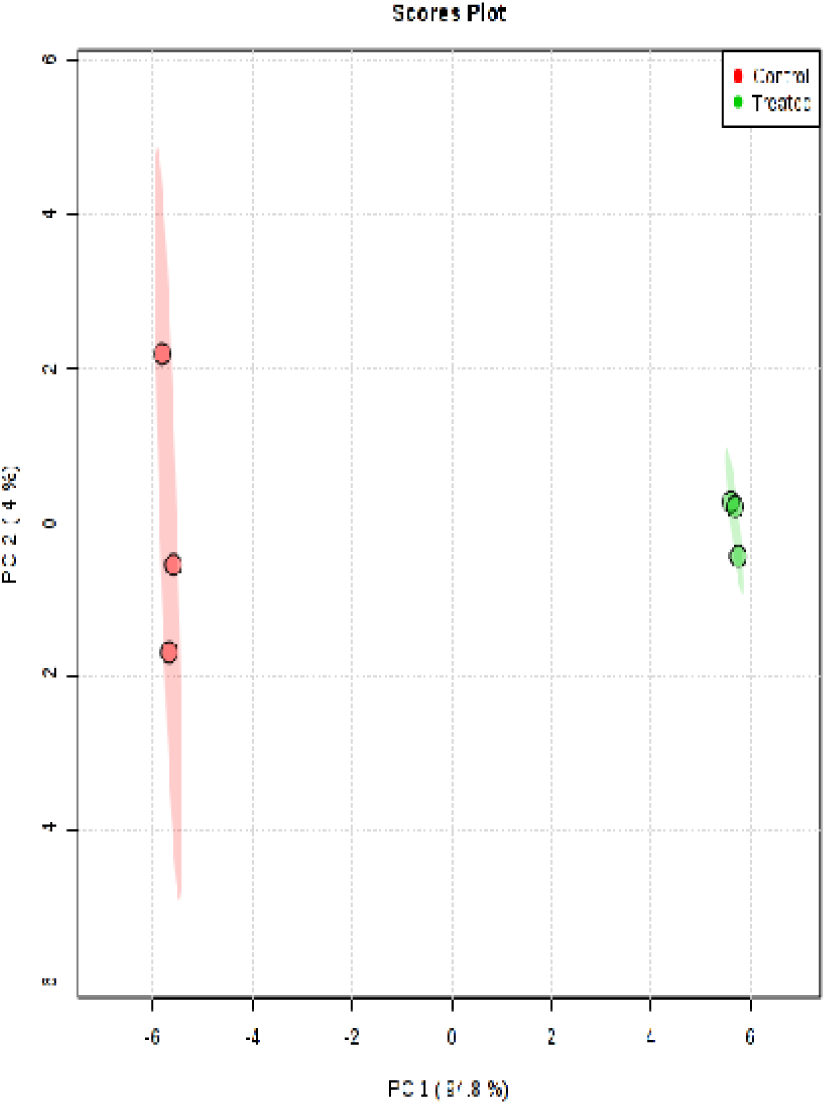
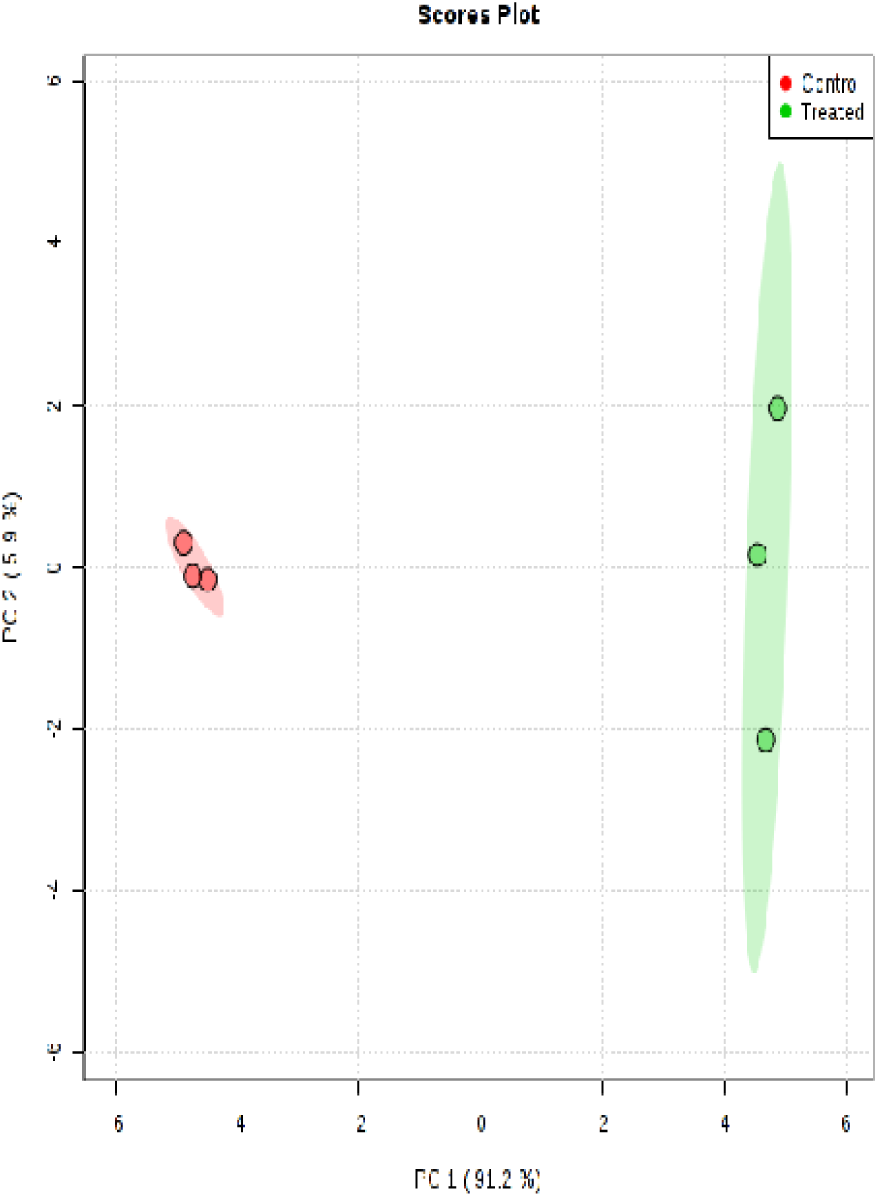
**a.** Score plot of tolerant roots. Control roots shows larger variation in metabolites compared to treated roots **b.** Score plot of sensitive roots. Control roots shows less diversity in metabolites compare to treated roots.

The combine data of tolerant genotype of leaf and roots shows higher fold change in octanoic acid, that are involved in fatty acid biosynthesis, 9-Decen-1-ol related to alcohol dehydrogenase and Erythrodiol an antioxidant. In roots, fold change was high of carboxylic acids (Pelargonic acid, Valeric acid, oxalic acid, sebacic acid, acetic acid) most of them have unpleasant odor might plays defensive role against various biotic stresses. Therefore it seems that metabolites of tolerant genotype are more uniform, shows coherent pattern. While, sensitive genotype leaves fold change was higher in Compounds like Iodononane, Nonyl Bromide, Docosane, Isophytol, Citronellyl, Tocopherol acetate, Carvon, Heptyl hexyl ether, 1-Napthalene acetic acid, Squalene, Phthalate. Most of them organic compound, aldehydes and terpenoids with unpleasant odor. Also Octodecanal (Stearaldehyde) which is pheromone. Whereas, roots of sensitive genotype contains Tiglic acid, Furan carboxylic acid, Valeric acid, Furan anhydride, Butyl glyoxalate, are carboxylic acids that are highly volatile and toxic secreted in response to various insects, bacteria. Caryophyllene is sesquiterpine, non-steroidal help to attracts insects. However, metabolites such as Nitropropane, 2-Nonane have mild fruity, sweet odor may help to attract insects that help in pollination. Hence sensitive genotypes in response to multiple stress release various metabolites with random pattern.

Comparative metabolomics analyses of two combined stresses response have highlighted a number of metabolites involved in diverse metabolic pathways. The Pathway analysis of the tolerant leaf and roots showed biosynthesis of unsaturated fatty acids, Glyoxalate and dicarboxylate metabolism, linoleic metabolism, fatty acid metabolism, Nicotinate and nicotinamide metabolism, Sulfur metabolism, pyruvate metabolism, and Glycolysis and gluconeogenesis metabolism. However, the pathway impact is highest of Nicotinate and Nicotinamide metabolism, sulfur metabolism and Pyruvate metabolism. Whereas, Sensitive genotype leaf and roots the number of pathways are more compared to tolerant genotype. Mainly the impact was high of the following pathways, Steroid biosynthesis, Sulfur metabolism, Panthothenate and Co-enzyme biosynthesis, Sesquiterpine and terpenoids biosynthesis, Arginine and Proline metabolism, Valine, leucine and isoleucine degradation, While, Aminoacyl t-RNA synthesis, Glucosinolate biosynthesis and Valine, Lucin and Isoleucine synthesis had no impact **(Supplemetary Table S1)**,.

### Heat map and correlation studies of metabolites

The heat map leaf and roots metabolites of tolerant genotype shows unique pattern. Although, the leaf metabolites of the tolerant genotypes differed with root metabolites. Almost all metabolites tolerant genotype leaf have higher concentration (=1), and some metabolites of treated leaf have concentrations above one (>1) like, Octanoic acid, 9-Decen1-ol, Erythediol. The concentration >1 of the following metabolites 1,16 dichloro hexadecane, 1-Pentanol, Eicosternoic acid, Lenolenic acid and ethyl Nonane. On the contrary, treated roots metabolites have very high concentrations the same metabolites in control roots shows v low concentration. Therefore the roots and leaf metabolites have no correlation with each other **Fig 3**.

**Fig 3.**
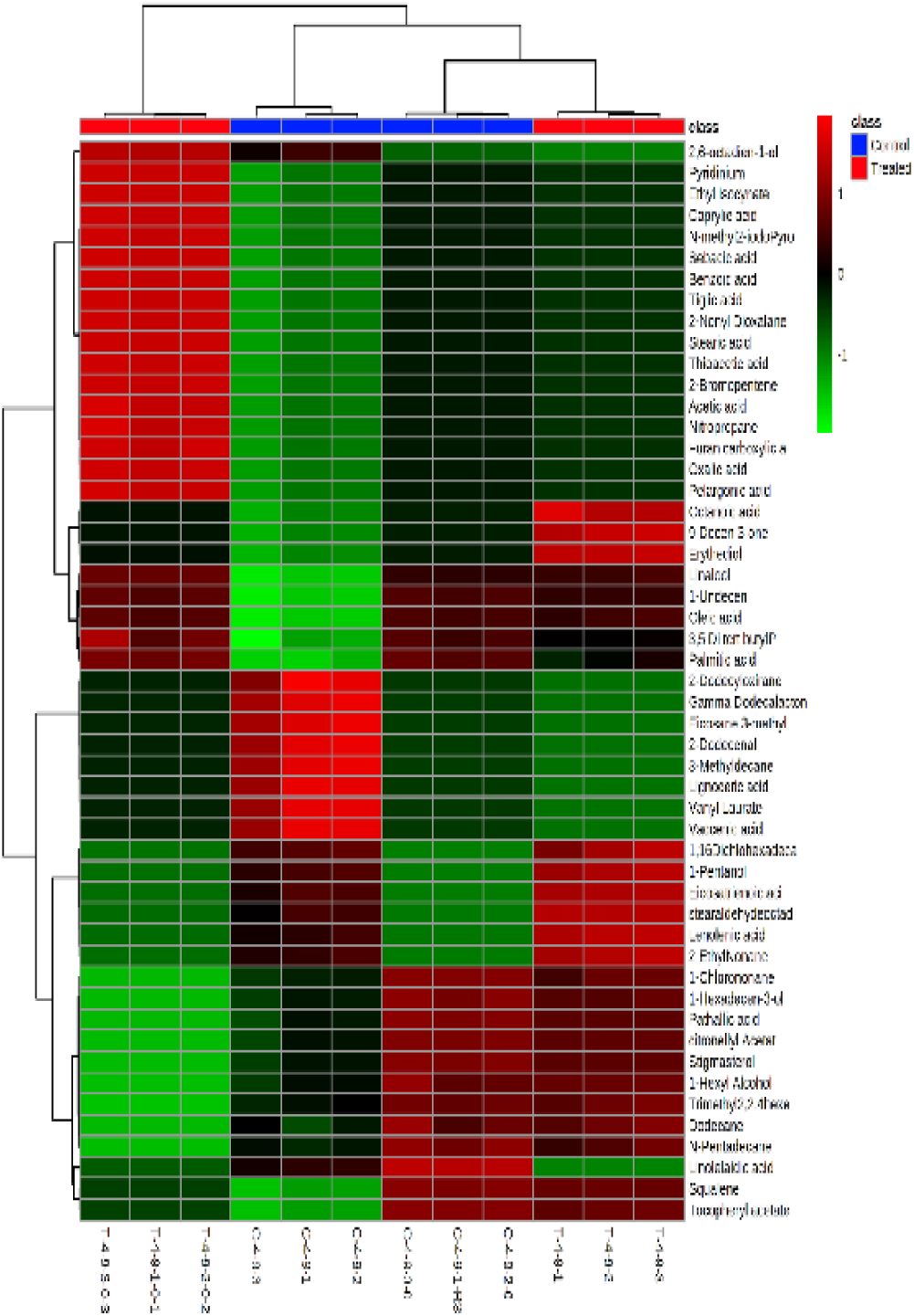
Heat map and Hierarchical clustering showing correlation between leaf and roots metabolites of tolerant genotype in response to combined abiotic stresses. The Red colors scale represents the high and Green - low concentrations of metabolites in control and treated plants.

Whereas, leaf and roots metabolites of sensitive genotype categorized into low and high concentrations **Fig 4**.

**Fig 4.**
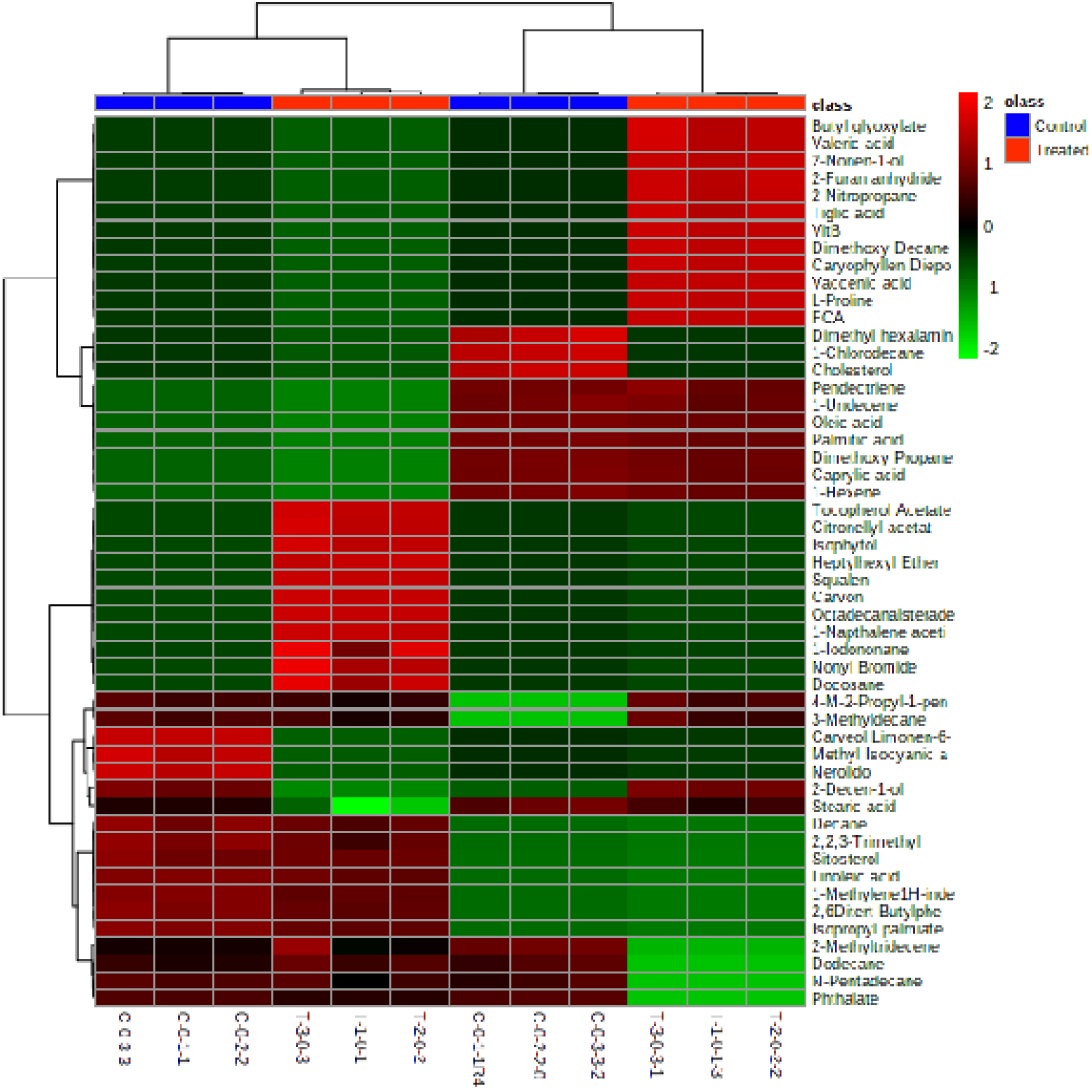
Heat map and Hierarchical clustering displays correlation between roots and leaf of sensitive genotype in response to combined abiotic stresses in control and treated. Red represent the –high concentration of metabolites and green represent the low concentration of metabolites.

Some of the metabolites of control roots are correlated with control and treated leaf (Dodecane, N-Pentadecane and Pthalate) shows higher concentration. Besides, Pentadectrine, 1-Undecen, oleic acid, palmatic acid, Dimethoxy propane, caprylic acid and 1-Hexene are common metabolites in control and treated roots with high concentrations. Therefore, in sensitive genotypes the leaf, roots metabolites are co-related or common. Thus there was less variations of metabolites in sensitive genotypes.

### Data analysis

The GC-MS data was analyzed using MetaboAnalyst (www.metaboanalyst.ca) web server. Peak areas of identified metabolites in triplicate in a form of comma separated excel file (.CVS format) were uploaded in this software. The software, check the data integrity of the uploaded files automatically. MetaboAnalyst replaces zero values with a small positive value (the half of the minimum positive number detected in the data). Data filtering, and data normalization was done by pooling controlled groups. The internal data structure was transformed to a table with each row represents a tissue sample (root, leaf) from tolerant genotype and sensitive genotypes and each column representing a feature (a metabolite). The data structure in this format, we performed a row-wise data normalization. The purpose of Row-wise normalization was to normalize each sample (row) so that they are comparable to each other. In Metaboanalyst, correlation analysis shows the correlation between different features. Further, hierarchical clustering analysis was done with the squared Euclidean distance and the average linkage method (between groups).The clustering result in the form of a heatmap. The values of the metabolites mean abundances were used to cluster the metabolomic data.

## 4. Discussion

### Role of tolerant genotype metabolites in multiple stresses

In the present work untargeted metabolites were quantified by GCMS technique and differentially regulated metabolites of tolerant (CML49) and sensitive (CML100) were compared with their respective controls. The major observation was differences in metabolite levels in treated versus untreated plants of both genotype, though less metabolites differed in treated leaves and roots of sensitive genotype comparative tolerant one respectively, these variations caused by simultaneous exposure of genotypes to various stresses and it depend on plants response also. The leaves metabolites of tolerant genotype shows fold increase in terpenoids linalool (monoterpen) and erythrodiol (triterpein) the main role of isoprenoid family have been associated in protection against oxidative and other abiotic stresses, especially volatile isoprenoids (Monoterpen) protective in photosynthetic processes during thermal and oxidative stresses (**Claudia Vikers et al. 2009**). Its role in membrane stabilization **(Sharkey and Singsaas 1995)** a hypothesis supports terpenes behave directly as an antioxidant, scavenges ROS by reactions through the conjugated double bond system **(Loretu and Velikova 2001; Affeck and Yakir 2002)**. Similarly, Erythrodiol is a pentacyclic triterpenes, leaf surface contains triterpenoids as constituents of waxes being involved in different roles such as maintenance of leaves structure, provide water, permeability, and plant insect interactions **(Bauer et al 2004; Mintz-Oron et al 2008; Wilson et al 2008)**. Correspondingly, non-volatile isoprenoid like Tocopherol acts as lipid soluble redox buffer and important scavenger of singlet oxygen species **(Foyer et al 2005)**. Also function as an antioxidants, in transgenic tobacco leaves its deficiency led to an enhancement in the lipid peroxidation **(Abbasi et al 2009)**. Majority of the metabolites accumulated in roots of tolerant genotype were belongs to saturated and unsaturated fatty acids, FAs and their derivatives directly involved in providing resistance against the bacterial (C16) and fungal pathogen (18:2 and 18:3) hence responsible for plants basal immunity **Kachroo and Kachroo (2009)**. Moreover, indirect and direct role of 18:0, 18:1, 18:2, and 18:3 plant defense signaling. In parsley, fungal infection rapidly induces the transcription of the 18:3-synthesizing omega-3 FA **(Kirsch et al 1997)**. Significant role of Oleic acid as a stimulator of the signaling enzyme phospholipase D (PLD) that function in anti-death cell **(Zhang et al. 2003)**. However, free fatty acids not only regulate abiotic stress tolerance, but also involve defenses against wound-induced responses and insect/herbivore feedings **(Upchurch 2008)**. In our studies, most of the metabolites of tolerant were involved in Phosphotidiyal and sphingolipid metabolism. Further work of **Yozo Okazaki and Kazuki Saito (2014)** confirms that membrane lipids has specific signal mediator and sensor of various biotic and abiotic stresses. Hence activating defense reactions at transcriptional level. As the lipids are stress mitigators, presence of polyunsaturated fatty acids that reduced cell damage expose to stress by scavenging reactive oxygen species. Therefore reducing the intensity of environmental stresses **(Mene-Saffrane et al 2009)**. Also, there are reports on membrane lipid remodeling in response to various stresses like, freezing, dehydration and nutrition depletion **(Moellering et al 2010; Gasulla et al 2013; Okazaki et al 2013b)**, and this remodeling contributes to the uphold membrane integrity and membrane-bound protein functions. While, Benzoic acid related to salicylic acid, (phenolic phytohormone) involve as an endogenous signal mediating plant defense responses against pathogens. Besides, it influence responses to abiotic stresses and related to aspects of plant growth and development **(Vlot et al 2009; Rivas-San Vicente and Plasencia 2011)**. The presence of haloalkanes (2-Bromopentene, ethyl isocynates) heterocyclic compound (2-Nonyl-1,3-dioxalane), hetrocyclic aromatic compounds (N-Methyl-2-Iodopyrrol), Methyl ester (Thioacetic acid) and Methyl ketone (3-Nitropropane) all these compound processes strong, unpleasant and odor may involve in repelling herbivory insects, pest and bacteria, fungi, nematodes (**Lamberth et al 2012; Francis et al 2013)**. The role of low molecular weight heterocyclic compounds derivatives of pyrimidine, pyrazole and isoflavones as a stimulators of seed germination and growth of maize seedlings **(Tsygankova et al 2016)**. Likewise, N-methyl-Δ-^1^Pyrrolinium is a cation combined with nicotinic acid to forms the alkaloid (S)-nicotine in roots of tobacco plant **(Katoh et al. 2005)** that inhibits a wide variety of herbivores **(Rosenthal G A 1986)**. *Trichoderma harzianum*, used as biocontrol agent, it produced Nonanoic acid (NA) that was very inhibitory to spore germination and mycelial growth of two cacao pathogens, Crinipellis perniciosa Stahel and Moniliophthora roreri Cif **(Aneja et al 2005)**. The antibiotic effect of acetic acid on some bacteria, including *Salmonella typhimurium, Escherichia coli* O157, and *Listeria monocytogenes* **(Delaquis et al 1999)**. The soluble oxalic acid is a strong acid, besides being toxic and lethal to grazing animals **(Von Burg 1994)**. It showed inhibitory effects on sucking insects’ planthopper and aphids **(Yoshihara et al 1980; Massonie 1980)**. Therefore we concluded that most of the roots metabolites were involved in providing stability to membranes and plant defense.

### Role of sensitive genotypes metabolites in multiple stresses

In sensitive leaf and root shows the accumulation of various metabolites, belongs to different classes, like terpenoids, Steroids, Ether lipids, Hydroquinone lipids, fatty aldehyde, fatty alcohol ester, fatty acyl conjugates, cyclic anhydrides, carboxylate, and linear alkanes. The leaves shows the emission of main terpenoids limnonen-oxide (carvon) and citronell. They are allelopathic in nature thus restricting seed germination and growth of nearby plants **(Maffei and Gertsch 2011)**. Moreover, plants emitted volatiles, also perceive or recognize them in inter- and intra-plant communications, so that plants under herbivore attack can alert neighbouring plant species, and priming their chemical defenses **(Heil 2014)**. The below ground emission of volatiles is an efficient trait: in maize roots the sesquiterpene (E)-β-caryophyllene is necessary to attract entomopathogenic nematodes to roots damaged by the ferocious maize pest Diabrotica virgifera. Maize varieties that lack this signal have been shown to be far more vulnerable to maize pest **(Rasmann et al 2005; Rasmann and Turlings 2007)**. However, there were some reports, that indicate due to lipophilic nature of monoterpenes the physical arrangement of phospholipids in the membranes were distorted and thus affect membrane functioning **(Zunino and Zygadlo 2004)**. Further, **Kaur et al (2010)** showed that citronellol generates ROS that induced lipid peroxidation, leakage of membrane and altered the membrane structure and permeability in wheat. Markedly as reported **Takshak and Agarwal (2015)** in UV-B treated leaf of *W somnifera* accumulated higher amount of phyto-constituents like Isophytol, trans-squalene, nonacosane and these plants were effective in scavenging free radicals. Furan fatty acids and their role in plants have been reviewed in detail **(Mowlong et al 2014)**. F-acids are mainly bound to phospholipids, they substitute for PUFAs and their strong capability (F-acids) to serve as radical scavengers suggests that plants use them to defend against oxidative stress **(Spiteller 2005)**. Moreover, the insecticidal activity of some 2-carboxylbenzofurans and their coumarin precursors against sweet potato weevil, *Cylas formicarius elegantulus* first demonstrated by **Williams et al (1994)** and later insecticidal properties of Benzofuran 2-carboxylic acid derivatives was reported **Jackson et al (1998)**. Water deficit led to produced various compounds in plants **Carmo-Silva et al (2009)** observe that in C4 grasses the amount of 5-HNV(2-amino-5-hydroxypentanoic acid (5-hydroxynorvaline, 5-HNV) in leaves of, C. dactylon and Z. japonica, increased under water deficit condition. However, any benefits from this unusual non-protein amino acid for drought resistance should be further explored to see that if it plays an important role in stress defensive mechanisms.

## 5 Conclusion

The metabolomics study of leaf and roots in response to combined abiotic stresses of two maize genotypes tolerant and sensitive showed the varied metabolic profile. Metabolomic profile of tolerant genotype have uniform pattern mostly involved in the biosynthesis of fatty acid metabolism, biosynthesis of saturating and unsaturated fatty acids, sulfur metabolism, thereby, regulated signaling, re-fix soil nitrate metabolism and defense against the wide variety of pathogens. Therefore, enhancing the tolerant mechanism of the genotype. In contrast, sensitive plant metabolites mostly involved in leaking or disrupting the membranes that disrupt signaling between various organelles and also for various stresses, defense related metabolites were accumulated in random pattern. either defense against the particular pathogens or pollination attracters. In present roots metabolomics study we can say, the metabolites somehow help to survive the sensitive genotype in multiple stress conditions but most of them are the causes of susceptibility. On the basis of the above observation, we can conclude that tolerant genotype plants by remodeling of the cell wall, maintain metabolic homeostasis, and proper signaling thus able to tolerate multiple stress conditions. Therefore, the study provides a comprehensive analysis of tolerance mechanism.

## Abbrevations

GC-MS: Gas chromatography mass spectrometry
Low-N: Low nitrogen stress
DAS: Days after sowing
PCA: Principal component analysis

## Acknowledgments

I acknowledge the grant received from DST Ministry of Science, Government of India, under the WOS-A scheme.

***Reference No: SR/WOS-A/LS-284/2013***

*“The authors have declared no conflict of interest”*.

## REFERNCES

Abbasi A. R, Saur A, Hennig P, Tschiersch H, Hajirezaei M, Hofius D, et al. (2009). Tocopherol deficiency in transgenic tobacco (*Nicotianatabacum* L.) plants leads to accelerated senescence. Plant Cell Environ 2009; 32: 144–157.

Affek, H.P. & Yakir, D. Protection by isoprene against singlet oxygen in leaves. Plant Physiol 2002; 129: 269–277.

Ai T N, Naing A H, Yun B-W, Lim S H and Kim C K. Overexpression of RsMYB1 Enhances Anthocyanin Accumulation and Heavy Metal Stress Tolerance in Transgenic Petunia. Front. Plant Sci 2018; 9: 1388.

Alfonso Albacete, Elena Cantero-Navarro, Dominik K, Großkinsky Cintia L, Arias María Encarnación, Balibrea Roque, Bru Lena Fragner, Michel E Ghanem, María de la Cruz, González Jose A, Hernández Cristina, MartínezAndújar Eric, van der Graaff, Wolfram Weckwerth, Günther Zellnig, Francisco Pérez-Alfocea, and Thomas Roitsch. Ectopic overexpression of the cell wall invertase gene CIN1 leads to dehydration avoidance in tomato. J Exp Botany 2015; 66 (3): 863–878

Alvarez, C., Garcia, I., Moreno, I., Perez-Perez, M.E., Crespo, J.L., Romero, L.C., and Gotor, C. Cysteine-generated sulfide in the cytosol negatively regulates autophagy and modulates the transcriptional profile in *Arabidopsis*. Plant Cell 2012b; 24: 4621–4634.

Aprile, A., Havlickova, L., Panna, R., Marè, C., Borrelli, G. M., Marone, D., et al. Different stress responsive strategies to drought and heat in two durum wheat cultivars with contrasting water use efficiency. BMC Genomics 2013; 14:821.

Bae M S, Cho E J, Choi E Y, and Park O K. Analysis of the Arabidopsis nuclear proteome and its response to cold stress. Plant Journal 2003; 36: 652–663.

Barbrook AC, Howe CJ, Purton S. Why are plastid genomes retained in non-photosynthetic organisms? Trends in Plant Science 2006; 11: 101–108.

Bauer, S., Schulte, E., and Thier, H.-P. (2004). Composition of the surface wax from tomatoes I. Identification of the components by GC/MS. European Food Research and Technology 2004; 219:

Bevan M. et al. 1998. Analysis of 1.9 Mb of contiguous sequence from chromosome 4 of *Arabidopsis thaliana*. Nature 1998; 391: 485–488.

Bermúdez, M.A., Galmés, J., Moreno, I., Mullineaux, P.M., Gotor, C., and Romero, L.C. Photosynthetic adaptation to length of day is dependent on S-sulfocysteine synthase activity in the thylakoid. Plant Physiol 2012; 160, 274–288.

Bradford M M. A rapid and sensitive method for the quantitation of microgram quantities of protein utilizing the principle of protein-dye binding. Anal Biochemistry 1976; 721 (2): 248–254.

Bui, Q. T., and Grandbastien, M. -A. (2012).“LT Rretrotransposons as controlling elements of genome response to stress?,”in Plant Transposable Elements: Impact on Genome Structure and Function Topics in Current Genetics, eds M.-A. Grandbastien and J.M. Casacuberta (Berlin;Heidelberg:Springer),273–296.

Cantoni G L. Biological methylation: selected aspects. Annu Rev Biochem 1975; 44: 435–451.

Ana E. Carmo-Silva, Alfred J. Keys, Michael H. Beale, Jane L. Ward, John M. Baker, Nathaniel D. Hawkins, Maria Celeste Arrabaça, Martin A.J. Parry. Drought stress increases the production of 5-hydroxynorvaline in two C4 grasses. Phytochemistry 2009; 70: 664–671.

Cecilia Gotor, Irene García, José L. Crespo & Luis C. Romero. Sulfide as a signaling molecule in autophagy, Autophagy 2013; 9: (4), 609–611.

Delaquis PJ, Sholberg PL, Stanich K (1999) Disinfection of mung bean seed with gaseous acetic acid. J Food Prot 1999; 62: 953–957.

Deserah D. Strand, Nicholas Fisher, and David M. Kramer. The higher plant plastid NAD (P)H dehydrogenase-like complex (NDH) is a high efficiency proton pump that increases ATP production by cyclic electron flow. J Biol Chem 2017; 292(28): 11850–11860.

Down’s M T. An improved hydrazine reduction method for the automated determination of low nitrate levels in freshwater. Water Res 1978; 12: 673–675.

Dubald M, Barakate A, Mandaron P, Mache R. The ubiquitous presence of exopolygalacturonase in maize suggests a fundamental cellular function for this enzyme. Plant J 1993; 4: 781–791.

Ebbs S, J Bushey, S Poston, D Kosma, M Samiotakis, D Dzombak. Transport and metabolism of free cyanide and iron cyanide complexes by willow. Plant Cell Environ 2003; 26: 1467–1478.

Essemine J, Qu M, Mi H and Zhu X-G. Response of Chloroplast NAD (P) H Dehydrogenase-Mediated Cyclic Electron Flow to a Shortage or Lack in Ferredoxin-Quinone Oxidoreductase-Dependent Pathway in Rice Following Short-Term Heat Stress. Front. Plant Sci 2016; 7:383.

Evans H.J., and Nason, A. Pyridine nucleotide-nitrate reductase from extracts of higher plants. Plant Physiol 1953; 28: 233–254. doi: 10.1104/pp.28.2.233.

Erice, G., S. Louahlia, J.J. Irigoyen, M. Sanchez-Diaz, J.C. Avice. 2010. Biomass partitioning, morphology and water status of four alfalfa genotypes submitted to progressive drought and subsequent recovery. Journal of Plant Physiology 167(2):114–120.

Foyer C H, and Noctor G. (2005) Oxidant and antioxidant signaling in plants: a re-evaluation of the concept of oxidative stress in a physiological context. Plant Cell Environ 2005; 29: 1056–1107.

Garcia, I., Castellano, J.M., Vioque, B., Solano, R., Gotor, C., and Romero, L.C. Mitochondrial {beta}-cyanoalanine synthase is essential for root hair formation in *Arabidopsis thaliana*. Plant Cell 2010; 22: 3268–3279.

GarcõÂa-Romera I, Fry S C. Upper limits for endogenous oligogalacturonides and free galacturonic acids in rose cell-suspension cultures: Implications for the action of exo- and endo-polygalacturonases in vivo. J Plant Physiol 1997; 150: 241±246.

Grandbastein M A. Activation of plant retrotransposons under stress conditions. Trends in Plant Sciences 1998; 3: 5 181–187.

Marie-Angèle Grandbastien, Hélène Lucas, Jean-Benôit More, Corinne Mhiri, Samantha Vernhettes & Josep M. Casacuberta. The expression of the tobacco Tnt1 retrotransposon is linked to plant defense responses. Genetica 1997; 100: 241–252.

Gotor, C., and Romero, L.C. S-sulfocysteine synthase function in sensing chloroplast redox status. Plant Signal. Behav 2013; 8: e23313.

Grover H L, Nair T V R, and Abrol Y P. Nitrogen metabolism of the upper three leaf blades of wheat at different soil Nitrogen levels. Physiol Plant 1978; 42: 287–292.

Guillas, I., Guellim, A., Reze, N. and Baudouin, E. Long chain base changes triggered by a short exposure of Arabidopsis to low temperature are altered by AHb1 non-symbiotic haemoglobin overexpression. Plant Physiol Biochem 2013; 63: 191–195.

Guo W, Zhao J, Li X, Qin L, Yan X, Liao H. A soybean b expansin gene GmEXPB2 intrinsically involved in root system architecture responses to abiotic stresses. Plant J 2011; 66(3):541–552.

Hageman, R.H., and Hucklesby, D.P. Nitrate reductase from higher plants. Meth. Enzymol 1971; 23: 491–503.

Hammani K, and Barkan A. An mTERF domain protein functions in group II intron splicing in maize chloroplasts. Nucleic Acids Research 2014; 42 (8): 5033–5042.

Han Y, Chen Y, Yin S, Zhang M, Wang W. Over-expression of TaEXPB23, a wheat expansin gene, improves oxidative stress tolerance in transgenic tobacco plants. J Plant Physiol 2015; 173:62–71.

Heil M. Herbivore-induced plant volatiles: targets, perception and unanswered question. New Phytologist 2014; 204: 297–306.

Hu, Xiuli; Liuji Wu, Feiyun Zhao, Dayong Zhang, NanaLi1 Guohui, Zhu Chaohao Li and Wei Wang. Phosphoproteomic analysis of the response of maize leaves to drought heat and their combination stress. Front Plant Sci 2015; 6: 1–21.

Iftekhar Alam, Dong-Gi Lee, Kyung-Hee Kim, Choong-Hoon park, Shamima Akhtar Sharmin, Hyoshin Lee Ki-Won OH Byung-Wook Yun and Byung-Hyun. Proteome analysis of soybean roots under waterlogging stress at an early vegetative stage. Journal of Bioscience 2010; 35: 1 49–62.

Ismond, K. P., Dolferus, R., dePauw, M., Dennis, E. S., and Good, A. G. Enhanced low oxygen survival in Arabidopsis through increased metabolic flux in the fermentative pathway. Plant Physiol 2013; 132: 1292–1302.

Ismond Aardra Kachroo and Pradeep Kachroo. 2009. Fatty acids-Derived signals in plant defense. Annu Rev. Phytopatho 17: 153–76.

Kaur K, Gupta AK, Kaur N. Effect of water deficit on carbohydrate status and enzymes of carbohydrate metabolism in seedlings of wheat cultivars. Indian Journal of Biochemistry and Biophysics 2007; 44: 223–230..

Kaur, Shalinder Shivani Ranab, Harminder Pal Singha,, Daizy Rani Batishb, and Ravinder Kumar Kohlib Citronellol Disrupts Membrane Integrity by Inducing Free Radical Generation. Z. Naturforsch 2011; 66 c: 260 – 266.

Kevin Francis, Crystal Smitherman, Shirley F. Nishino, Jim C. Spain, Giovanni Gadda. The Biochemistry of the Metabolic Poison Propionate 3-Nitronate and Its Conjugate Acid, 3-Nitropropionate. International Union of Biochemistry and Molecular Biology 2013; 65: (9): 759–768.

S. Kimura, K. Sakaguchi, DNA repair in plants, Chem. Rev. 2006; 106: 753–766.

Kirsch C, Takamiya-Wik M, Reinold S, Hahlbrock K, Somssich IE. Rapid, transient, and highly localized induction of plastidial omega-3 fatty acid desaturase mRNA at fungal infection sites in *Petroselinum crispum*. Proc. Natl. Acad. Sci. USA 1997; 94:2079–84.

Klára Kosová, Pavel Vítámvás, Milan O Urban, Ilja T Prášil and Jenny Renaut. Plant Abiotic Stress Proteomics The Major Factors Determining Alterations in Cellular Proteome. Frontiers in Plant Science 2018; 122: 9 1-22–322.

Komatsu S Deschamps, Thibaut Susumu Hiraga, Mikio Kato Mitsuru, Chiba Akiko Hashiguchi, Makoto Tougou Satoshi, Shimamura Hiroshi Yasue. Characterization of a novel flooding stress-responsive alcohol dehydrogenase expressed in soybean roots. Plant Molecular Biology 2011; 77: 309.

Kun Zhang, Huiting Cui, Shihao Cao, Li Yan Mingna Li1, Yan Sun. 2019. Overexpression of *CrCOMT* from *Carex rigescens* increases salt stress and modulates melatonin synthesis in *Arabidopsis thaliana*. Plant Cell Reports 223–228.

Kwasniewski M, Szarejko I. Molecular cloning and characterization of beta-expansin gene related to root hair formation in barley. Plant Physiol 2006; 141(3):1149–1158.

Jackson Y A, Mark F. Williams, Lawrence A. D. Williams, Kisha Morgan & Flona A. Redway. Insecticidal Properties of Benzofuran-2-carboxylic Acid Derivatives. Pestic. Sci 1998; 53: 241–244.

Jander G, Joshi V. Recent progress in deciphering the biosynthesis of aspartate-derived amino acids in plants. Mol Plant 2010; 3:54–65.

Lang, Nguyen Thi; Nguyen Quang Cao Binh, Chau Thanh Nha, Bui Chi Buu. A candidate gene response to drought stress condition in rice (Oryza sativa L.). Omonrice 2010; 17: 105–113.

Lee, S.; Lee, E.J.; Yang, E.J.; Lee, J.E.; Park, A.R.; Song, W.H.; Park, O.K Proteomic identification of annexins, calcium-dependent membrane binding proteins that mediate osmotic stress and abscisic acid signal transduction in Arabidopsis. Plant Cell 2004; 16: 1378–1391.

Lee, Y. and Kende, H. Expression of a-expansin and expansin-like genes in deepwater rice. Plant Physiol 2002. 130: 1396–1405.

Li AX, Han YY, Wang X, Chen YH, Zhao MR, Zhou S-M, Wang W (2015a) Root-specific expression of wheat expansin gene TaEXPB23 enhances root growth and water stress tolerance in tobacco. Environ Exp Bot 2015; 110:73–84.

Lianwei Peng, Hiroshi Yamamoto, Toshiharu Shikanai. Structure and biogenesis of the chloroplast NAD(P)H dehydrogenase complex. Biochimica et Biophysica Acta 2011; 1807: 945–953.

Lisch D. How important are transposons for plant evolution? Nat. Rev. Genet 2012; 14: 49–61.

Liu H, Ma Y, Chen NA, Guo S, Liu H, Guo X, Chong K, Xu, Y. Overexpression of stress-inducible OsBURP16, the β subunit of polygalacturonase 1, decreases pectin content and cell adhesion and increases abiotic stress sensitivity in rice. Plant Cell and Environment 2013; 37: 1144–1158.

Loreto, F., Forster, A., Durr, M., Csiky, O. & Seufert, G. On the monoterpene emission under heat stress and on the increased thermotolerance of leaves of Quercusilex L. fumigated with selected monoterpenes. Plant Cell Environ 1998; 21: 101–107.

Loreto, F and Velikova, V. Isoprene produced by leaves protects the photosynthetic apparatus against ozone damage, quenches ozone products, and reduces lipid peroxidation of cellular membranes. Plant Physiol 2001; 127: 1781–1787.

Clemens Lamberth, Stephan Trah, Sebastian Wendeborn, Raphael Dumeunier, Mikael Courbot, Jeremy Godwin, Peter Schneiter. Synthesis and fungicidal activity of tubulin polymerisation promoters. Part 2: Pyridazines. Bioorganic & Medicinal Chemistry 2012; 20: 2803–2810.

Lunn J E. Sucrose-phosphatase gene families in plants. Gene 2003; 303: 187–196.

Luis C. Romero, M. Ángeles Aroca, Ana M. Laureano-Marín, Inmaculada Moreno, Irene García, and Cecilia Gotor. Cysteine and Cysteine-Related signaling Pathways in *Arabidopsis thaliana*. Molecular Plant 2014; 7 (4): 264–276.

McClintock, B. The significance of responses of the genome to challenge. Science 1984; 226: 792–80.

Madhu Anejaa, Thomas J. Gianfagnaa, Prakash K. Hebbarb. Trichoderma harzianum produces nonanoic acid, an inhibitor of spore germination and mycelial growth of two cacao pathogens. Physiological and Molecular Plant Pathology 2005; 67: 304–307.

Ana Maldonado M, Sira Echevarría-Zomeño, Sylvia Jean-Baptiste, Martha Hernández, Jesús Jorrín-Novo V. Evaluation of three different protocols of protein extraction for rabidopsis thaliana leaf proteome analysis by two dimensional electrophoresis. J. Proteomics 2008; 71(4): 461–47.

Mao, H., Wang, H., Liu, S., Li, Z., Yang, X., Yan, J et al. A transposable elementina NAC gene is associated with drought tolerance in maize seedlings. Nat.Commun 2015; 6: 8326.

Massonie G (1980) Breeding of a biotype of Myzus persicae Sulzer on a synthetic medium. V. Influence of oxalic and gentisic acids on the nutritive value of a synthetic medium. Ann Nutr Aliment 1980; 34: 139–146.

Mawlong Ibandalin, M. S. Sujith Kumar, Dhiraj Singh. Furan fatty acids: their role in plant systems. Phytochem Rev 2014; 1–7.

Mène-Saffrané L, Dubugnon L, Chételat A, Stolz S, Gouhier-Darimont C, Farmer EE. Nonenzymatic oxidation of trienoic fatty acids contributes to reactive oxygen species management in *Arabidopsis*. J Biol Chem 2009;. 284:1702–708.

Mintz-Oron, S., Mandel, T., Rogachev, I., Feldberg, L., Lotan, O., Yativ, M., Aharoni, A. (2008). Gene expression and metabolism in tomato fruit surface tissues. Plant Physiology 2008; 147: 823–851.

Mittler, R. Abiotic stress, the field environment and stress combination. Trends Plant Sci 2006; 11: 15–19.

Moellering, E.R., Muthan, B. and Benning, C. Freezing tolerance in plants requires lipid remodeling at the outer chloroplast membrane. Science 2010; 330: 226–228.

Narendra Tuteja, Parvaiz Ahmad, Brahma B. Panda, Renu Tuteja (2009) Genotoxic stress in plants: Shedding light on DNA damage, repair and DNA repair helicases. Mutation Research 2009; 681: 134–149.

Okazaki, Y., Otsuki, H., Narisawa, T., Kobayashi, M., Sawai, S., Kamide, Y., Kusano, M., Aoki, T., Hirai, M.Y. and Saito, K. A new class of plant lipid is essential for protection against phosphorus depletion. Nat. Commun 2013b; 4: 1510.

Prasad, P. V. V., Pisipati, S. R., Momcilovic, I., and Ristic, Z. Independent and combined effects of high temperature and drought stress during grain filling on plant yield and chloroplast EF-Tu expression in spring wheat. J. Agron. Crop Sci 2011; 197: 430–441.

Piotrowski, M. Primary or secondary? Versatile nitrilases in plant metabolism. Phytochemistry 2008; 69: 2655–2667.

Ralph Reimholz, Peter Geigenberger, Mark Stitt. Sucrose-phosphate synthase is regulated via metabolites phosphorylation in potato tubers, in a manner analogous to the enzyme in leaves. Planta 1994; 192: 480–488.

Rasmann S. et al. Recruitment of entomo-pathogenic nematodes by insect-damaged maize roots. Nature 2005; 434: 732–737.

Rasmann, S., Turlings, T.C.J. Simultaneous feeding by aboveground and belowground herbivores attenuates plant-mediated attraction of their respective natural enemies. Ecology Letters 2007; 10: 926–936.

Reinhard Korbinian Proels and Ralph Huckelhoven. Cell-wall invertases, key enzymes in the modulation of plant metabolism during defence responses. Molecular Plant pathology 2014; 15(8): 858–864.

Rivas-San Vicente, M., and Plasencia, J. Salicylic acid beyond defence: its role in plant growth and development. J Exp.Bot 2011; 62: 3321–3338.

Rizhsk, L., Liang, H., Shuman, J., Shulaev, V., Davletova, S., and Mittler, R. (2004).When defense pathways collide. The response of Arabidopsis to a combination of drought and heat stress. Plant Physiol 2004; 134: 1683–1696.

Robles P, Micol JL, Quesada V. Arabidopsis MDA1, a Nuclear-Encoded Protein, Functions in Chloroplast Development and Abiotic Stress Responses. PLoS ONE 2012; 7(8): e42924.

Roitsch T, Balibrea ME, Hofmann M, Proels R, Sinha A K. Extracellular invertase: key metabolic enzyme and PR protein. J Exp Bot 2003; 54: 513–524.

Romero, L.C., Garcia, I., and Gotor, C. L-Cysteine Desulfhydrase 1 modulates the generation of the signaling molecule sulfide in plant cytosol. Plant Signal Behav 2013; 8: e24007

Rosenthal G. A. Scientific American 1986; 254: 76.

Rufty T W, Huber S C, Volk R J. Alternations in leaf carbohydrate metabolism in response to nitrogen stress. Plant Physiol 1988; 88:725–730.

Salekdeh GH, Siopongco J, Wade LJ, Ghareyazie B, Bennett J. Proteomic analysis of rice leaves during drought stress and recovery. Proteomics 2002; 2: 1131–1145.

Siegien I and R Bogatek. Cyanide action in plants: from toxic to regulatory. Acta Physiol Plant 2006; 28: 483–497.

Saori Ogawa and Shiro Mitsuya. *S*-methylmethionine is involved in the salinity tolerance of *Arabidopsis thaliana* plants at germination and early growth stages. Physiologia Plantarum 2012; 144: 13–19. 2012.

Sanchez-Perez R, Jorgensen K, Olsen C E, Dicenta F, Moller B L. Bitterness in almonds. Plant Physiol 2008; 146(3):1040–1052.

Shapiro J. A 21st century view of evolution: genome system architecture, repetitive DNA, and natural genetic engineering. Gene 2005; 345: 91–100.

Sharkey, T D and Singsaas, E L. Why plants emit isoprene. Nature 1995; 374: 769.

Solís-Guzmán, Marí.Gloria., Argüello-Astorga, G., López-Bucio, José., Ruiz-Herrera, Leó.Francisco., López-Meza, J.E., Sánchez-Calderón, L., Carreón-Abud, Yazmí., Martínez-Trujillo, M., *Arabidopsis thaliana* sucrose phosphate synthase (*sps*) genes are expressed differentially in organs and tissues, and their transcription is regulated by osmotic stress, Gene Expression Patterns 2017;

Spiteller G. Furan fatty acids: occurrence, synthesis, and reactions. Are furan fatty acids responsible for the cardioprotective effects of a fish diet?Lipids 2005; 40: 755–771.

Suzuki, N., Rivero, R. M., Shulaev, V., Blumwald, E., and Mittler, R. Abiotic and biotic stress combinations. New Phytol 2014; 203: 32–43.

Swidzinski JA, et al. A custom microarray analysis of gene expression during programmed cell death in Arabidopsis thaliana. Plant J 2002; 30: 431–446.

Tahir, J., Watanabe, M., Jing, H.C., Hunter, D.A., Tohge, T., Nunes-Nesi, A., Brotman, Y., Fernie, A.R., Hoefgen, R., and Dijkwel, P.P. Activation of R-mediated innate immunity and disease susceptibility is affected by mutations in a cytosolic O-acetylserine (thiol) lyase in *Arabidopsis*. Plant J 2013; 73: 118–130.

Takshak Swabha; S. B. Agrawal. Alterations in metabolite profile and free radical scavenging activities of Withania somnifera leaf and root extracts under supplemental ultraviolet-B radiation. Acta Physiol Plant 2015; 37:260.

Taylor N L, et al. Analysis of the rice mitochondrial carrier family reveals anaerobic accumulation of a basic amino acid carrier involved in arginine metabolism during seed germination. Plant Physiology 2010; 154: 691–704.

Tsygankova Victoria, Yaroslav Andrusevich, Olexandra Shtompel, Artem Hurenko, Roman Solomyannyj, Galyna Mrug, Mikhaylo Frasinyuk, Volodymyr Brovarets. Stimulating effect of five and six-membered heterocyclic compounds on seed germination and vegetative growth of maize (*Zea mays* L.) International Journal of Biology Research 2016; 1 (4): 01–14

Turner, M. F., Heuberger, A. L., Kirkwood, J. S., Collins, C. C., Wolfrum, E. J., Broeckling, C. D., et al. Non-targeted metabolomics in diverse Sorghum breeding lines indicates primary and secondary metabolite profiles are associated with plant biomass accumulation and photosynthesis. Front. Plant Sci 2016; 7: 953.

Upchurch R G. Fatty acid unsaturation, mobilization, and regulation in the response of plants to stress. Biotechnol. Lett 2008; 30:967–77.

Urbonavieius J, Qian Q, Durand J M B, Hagervall T G, and Bjork G R.. Improvement of reading frame maintenance is a common function for several tRNA modification. EMBO J 2001; 20: 4863–4873.

Velikova, V. and Loreto, F. On the relationship between isoprene emission and thermotolerance in Phragmites australis leaves exposed to high temperatures and during the recovery from a heat stress. Plant Cell Environ 2005; 28: 318–327.

Vile, D., Pervent, M., Belluau, M., Vasseur F.,Bresson, J., Muller, B., et al. Arabidopsis growth under prolonged high temperature and water deficit: independent or interactive effects? Plant Cell Environ 2012; 35: 702–718.

Vlot, A.C., Dempsey, D.M.A., and Klessig, D.F. Salicylic acid, a multi-faceted hormone to combat disease. Annu.Rev.Phytopathol 2009; 47: 177–206.

Von Burg R. Toxicology update. J Appl Toxicol 1994; 14: 233–237.

Wen, W., Li, K., Alseekh, S., Omranian, N., Zhao, L., Zhou, Y., et al. Genetic determinants of the network of primary metabolism and their relationships to plant performance in a maize recombinant inbred line population. Plant Cell 2015; 27: 1839–1856.

Wataru Yamori, Naoki Sakata, Yuji Suzuki, Toshiharu Shikanai and Amane Makino. Cyclic electron flow around photosystem I via chloroplast NAD (P)H dehydrogenase (NDH) complex performs a significant physiological role during photosynthesis and plant growth at low temperature in rice. The Plant Journal 2011; 68: 966–976.

Williams, L. A. D., Anderson, M. J. and Jackson, Y. A., Insecticidal activity of synthetic 2-carboxylbenzofurans and their coumarin precursors. Pestic Sci 1994; 42: 167–71.

Wilson, Shan, H., W. K., Phillips, D. R., Bartel, B., and Matsuda, S. P. T. Trinorlupeol: A major nonsterol triterpenoid in arabidopsis. Organic Letters 2008; 10: 1897–1900.

Wirtz, M., and Hell, R. Functional analysis of the cysteine synthase protein complex from plants: structural, biochemical and regulatory properties. J. Plant Physiol 2006; 163: 273–286.

Winkel-Shirley B. Flavonoid biosynthesis, A colorful model for genetics, biochemistry, cell biology and biotechnology. Plant Physiol 2001; 26:485–93.

Won S, Choi S, Kumari S, Cho M, Lee SH, Cho H. Root hair specific EXPANSIN B genes have been selected for Graminaceae root hairs. Mol Cells 2010. 30(4):369–376.

Xing SC, Li F, Guo QF, Liu DR, Zhao XX, Wang W. The involvement of an expansin geneTaEXPB23 from wheat in regulating plant cell growth. Biol Plant 2009; 53(3):429–434.

Yan J X, Wait R Berkelman, T Harry R A, et al. Electrophoresis 2000; 21: 3666–3672.

Yaeno T, Matsuda O, Iba K. Role of chloroplast trienoic fatty acids in plant disease defense responses. Plant J 2004; 40: 931–41.

Yip W K, Yang S F. Cyanide metabolism in relation to ethylene production in plant tissues. Plant Physiol 1988; 88(2):473–476.

Yoshihara T, Sogawa K, Pathak MD, Juliano BO, Sakamura S. Oxalic acid as a sucking inhibitor of the brown planthopper in rice (Delphacidae, Homoptera). Entomol Exp Appl 1980; 27: 149–155.

Yozo Okazaki and Kazuki Saito. Roles of lipids as signaling molecules and mitigators during stress response in plants. The Plant Journal 2014; 79: 584–596.

Zhao Y, Cai M, Zhang X, Li Y, Zhang J, et al. Genome-Wide Identification, Evolution and Expression Analysis of mTERF Gene Family in Maize. PLoS ONE 2014; 9(4): e94126.

Zhao D, Reddy K R, Kakani V G, Reddy V R. Nitrogen deficiency effects on plant growth, leaf photosynthesis, and hyperspectral reflectance properties of sorghum. Eur J Agron 2005; 22: 391–403.

Zhao, M.R., Han, Y.Y., Feng, Y.N., Li, F. and Wang, W. Expansins are involved in cell growth mediated by abscisic acid and indole-3-acetic acid under drought stress in wheat. Plant Cell Rep 2012. 31: 671–685.

Zhang W, Wang C, Qin C, Wood T, Olafsdottir G, Welti R, and Wang X. The oleate-stimulated phospholipase D, PLD delta, and phosphatidic acid decrease H2O2-induced cell death in Arabidopsis. Plant Cell 2003, 15: 2285–2295.

Zivy, M., Wienkoop, S., Renaut, J., Pinheiro, C., Goulas, E., and Carpentier, S. The quest for tolerant varieties: the importance of integrating “omics” techniques to phenotyping. Front. Plant Sci 2015; 6: 448.

Zou H, Wenwen Y, Zang G, Kang Z, Zhang Z, Huang J, Wang G. OsEXPB2, a b-expansin gene, is involved in rice root system architecture. Mol Breed 2015; 35(1):

Zunino M. P. and Zygadlo J. A. Effect of monoterpenes on lipid peroxidation in maize. Planta 2004; 219: 303 – 309.

